# Discovering cancer stem-like cells using Spatial transcriptomic analysis: Nuclear factor I X as a novel therapeutic target for gastric cancer

**DOI:** 10.1101/2024.03.31.587468

**Authors:** Akira Ishikawa, Takafumi Fukui, Aya Kido, Narutaka Katsuya, Kazuya Kuraoka, Naohiro Uraoka, Takahisa Suzuki, Shiro Oka, Takahiro Kotachi, Hassan Ashktorab, Duane Smoot, Wataru Yasui

## Abstract

**Background:** Gastric cancer (GC) is characterized by significant intratumoral heterogeneity and stem cells presenting as promising therapeutic targets. Despite advancements in spatial transcriptome analysis, unexplored targets for addressing cancer stemness remain unknown. This study aims to identify Nuclear Factor IX (NFIX) as a critical regulator of cancer stemness in GC and evaluate its clinicopathological significance and function.

**Methods:** Spatial transcriptome analysis was conducted on GC. NFIX expression’s correlation with clinicopathological factors and prognosis was assessed through immunostaining in 127 GC cases. Functional analyses in cancer cell lines validated these findings.

**Results:** Spatial transcriptome analysis stratified GC tissues based on genetic profiles, pinpointing CSC-like cells and further refined the classification to identify and highlight the NFIX’s significance, validated by Monocle 3 and CytoTRACE analyses. Knockdown experiments in cancer cell lines demonstrated the involvement of NFIX in cancer cell proliferation and kinase activity.

**Conclusions:** This study underscores spatial transcriptome analysis’s role in refining GC tissue classification and identifying therapeutic targets, highlighting NFIX as pivotal. NFIX expression correlates with poor prognosis and drives GC progression, suggesting its potential as a novel therapeutic target for personalized GC therapies.

## INTRODUCTION

Gastric cancer (GC), the third leading cause of cancer-related mortality in Japan (1), presents a substantial clinical challenge, particularly in advanced stages, where the prognosis markedly worsens (2). The complexity of GC pathogenesis is further compounded by the pivotal role of cancer stem cells (CSCs) (3), which drive tumor progression, metastasis, and therapeutic resistance (4). Previous studies on cancer stemness have highlighted molecules such as KIFC1 and Annexin A10 (ANXA10), especially in the stomach (9, 10), emphasizing the demand for novel therapeutic targets. Recent advancements in molecular biology and genomics have opened new avenues for targeted therapies; however, the identification and validation of effective targets in GC remain a critical area of ongoing research. The development of such targets could revolutionize treatment paradigms and significantly improve patient outcomes.

Spatial transcriptome analysis is currently attracting attention because of its potential to overcome intratumoral heterogeneity (11, 12). The 10x Genomics platform has revolutionized spatial transcriptomics by simplifying the transfer of transcriptomic probes from standard glass slides to Visium slides. The CytAssist system has been at the forefront of technological discussions in spatial transcriptomics. Its role in enhancing in situ hybridization techniques is pivotal for visualizing RNA molecules within their native tissue environments. CytAssist has expanded the scope of spatial transcriptomics, allowing for more sensitive and efficient gene expression analysis. This advancement enables a deeper understanding of tissue architecture and function (13, 14). Indeed, there is one report that examined the stomach using spatial analysis (15, 16); however, there are still no reports on GC stemness.

In this study, we identified CSC-like cells using spatial transcriptomic analysis. Among the distinctive genes characterizing CSC-like cells, Nuclear Factor IX (NFIX) has emerged as a gene of particular interest. Functional analyses of NFIX revealed significant clinical and pathological implications. This discovery not only enhances our understanding of GC pathogenesis but also opens new avenues for potential therapeutic interventions targeting CSC-like cells.

## MATERIALS AND METHODS

### Spatial Transcriptomics (CytAssist Visium)

Spatial transcriptomic construction and sequencing utilized formalin-fixed, paraffin-embedded (FFPE) samples that met RNA quality control criteria (DV200 > 50%).

Tissues were prepared according to the Visium CytAssist Spatial Gene Expression for FFPE Tissue Preparation Guide (10× Genomics, Pleasanton, CA, USA). Sequencing was performed at the Research Institute for Microbial Diseases, Osaka University. Libraries were sequenced using an MGI DNBSEQ-G400RS (MGI Tech Co., Shenzhen, China). The Space Ranger pipeline v2022.0705.1 (10× Genomics, Pleasanton, CA, USA) and the GRCh38-2020- A reference were used to process the FASTQ files.

Gene-spot matrices from the samples were analyzed using the Seurat R package (version 4.3.1) (17). For each data sample, spots were filtered to obtain a minimum detected gene count of 100 genes. Normalization across spots was performed using the SCTransform2 function. Dimensionality reduction and clustering were performed using independent component analysis. The correlation matrix of the spatial cluster genes was generated by initially selecting differentially expressed genes (average log fold change (logFC) > 0.25, adjusted p-value < 0.05, by Wilcoxon rank-sum test). The Pearson’s correlation coefficient was calculated across the matrix. Spatial feature expression plots were generated using the spatial feature plot function in Seurat (version 4.3.0).

Transcriptional entropy was calculated using CytoTRACE software (18). The computational analysis of compartment embedding trajectories was performed using the Monocle 3 algorithm, following the methodology outlined by Qiu et al. (19). This process involved transferring dimensionality-reduced data from the integrated and batch-corrected pools corresponding to each compartment to the cell_data_set objects within the Monocle 3 framework. For each individual compartment, the initial point of the trajectory was determined by identifying the node closest to the recognized progenitor population. From this initial node, the trajectory pseudotime is computed, providing a structured temporal framework for the analysis.

### Biological informatics analysis for RNA sequence

To explore the biological function of a cluster, we compared its expression value with the average value of all clusters within the same sample. We then obtained the genetic FC and ranked the gene list according to log2FC. We used the clusterProfiler R package to complete the gene set enrichment analysis, and the results were visualized using the ggplot2 R package, as described in our previous report (7). Canonical pathways were used as background gene sets for the pathway enrichment analysis. Molecular Function (MF) analysis, including spatial transcriptomic cluster 0 analysis and NFIX knockdown cell line analysis, utilized Gene Ontology gene sets.

### Tissue sample for Spatial Transcriptomics

A 73-year-old male with no family history of hereditary disease or cancer presented to our hospital and underwent endoscopic submucosal dissection of a gastric tumor. The tumor measured 13 mm in diameter and was located on the posterior wall of the greater curvature of the stomach. The pathological examination identified the carcinoma primarily as a well-differentiated tubular adenocarcinoma, with portions showing charateristics of non-solid-type, poorly differentiated adenocarcinoma. The tumor was staged as pT1a or cN0M0, indicating confinement to the mucosa (T1a) without regional lymph node involvement (cN0) or distant metastasis (cM0). Importantly, there was no evidence of vascular invasion or tumor exposure at the resection margins.

### Data availability

Raw and processed data reported in this paper were deposited in the Gene Expression Omnibus with accession code GSE245704 for spatial transcriptomic analysis and in the DNA Data Bank of Japan (DDBJ) with accession code PRJDB16815 for RNA sequencing.

### Tissue samples for immunohistochemistry and in situ hybridization

In this study, 127 primary GC samples were collected from patients who underwent curative resection between 2012 and 2015 at Kure Medical Center and Chugoku Cancer Center (Kure, Hiroshima, Japan), as previously described (7). Archived FFPE tumor tissues obtained from resected specimens were used for immunohistochemical analyses. One representative tumor block from each specimen underwent assessment by IHC. All GC tumor stages were based on the Japanese classification of gastric cancer and the tumor, node, and metastasis classification of the Union for International Cancer Control (20). For Epstein–Barr virus (EBV), all slides were stained using an automated slide staining system (Benchmark XT; Ventana Medical Systems, Inc., Tucson, AZ, USA). A ready-to-use EBER (EBV-encoded small RNA) probe (Ventana/Roche) was used together with ISH-Protease 3 pretreatment for 1 h and probe incubation for 28 min. Informed consent was obtained from all the patients. This study was approved by the Ethics Committee for Human Genome Research of Hiroshima University (E2001-9923), and was conducted following the guidelines of the Declaration of Helsinki.

### Molecular subtypes using surrogate markers

We divided 127 GC cases into four molecular subtypes according to a previous report(21). The microsatellite-instable (MSI) subtype was defined as tumors showing a loss or reduction in MLH1 of more than 30%. Strong nuclear staining of p53 (more than 50%) was used as a surrogate for the chromosomal instability (CIN) subtype. The remaining cases showing a loss or reduction of E-cadherin expression were categorized as having genomic stability (GS). The Epstein-Barr virus (EBV) subtype indicated EBER-positive tumors.

### Cell lines

The human GC-derived cell lines, MKN-1, MKN-7, MKN-45, and MKN-74, were purchased from the Japanese Collection of Research Bioresources Cell Bank (Osaka, Japan). These cell lines were cultured at 37 °C in Roswell Park Memorial Institute (RPMI)-1640 medium (#05911; Nissui Pharmaceutical, Tokyo, Japan) supplemented with 10% fetal bovine serum (#35-015-CV, Corning; Corning, NY, USA) in a humidified atmosphere with 5% CO2, following previously described protocols (22). The non-tumorigenic gastric cell line HFE-145, which was provided by H. Ashktorab and D. Smoot (Howard University, Washington, DC, USA) (23), was maintained at 37°C in Dulbecco’s modified Eagle’s medium (DMEM-F12) (#11320033, Thermo Fisher Scientific) containing 10% fetal bovine serum (#35-015-CV, Corning; Corning, NY, USA) in a humidified atmosphere with 5% CO2.

### IHC

For IHC, representative FFPE slides were cut into small sections (4 µm), deparaffinized, and rehydrated. Immunohistochemical analysis was conducted using a Dako EnVision+ Peroxidase Detection System (#K4003, Dako Cytomation, Carpinteria, CA, USA).

Briefly, heat-mediated antigen retrieval in FFPE tissue sections was performed in sodium citrate buffer (pH 9.0) for 50 min at 95°C. Peroxidase activity was blocked with 3% H2O2-methanol for 10 min. The sections were incubated with rabbit polyclonal anti-NFIX antibody (NBP2-15039, 1:200; Novus Biologicals, LLC, USA) for 1 h at room temperature, followed by incubation with EnVision+ peroxidase-conjugated anti-rabbit secondary antibody for 1 h. For the color reaction, the sections were incubated with the Dako Liquid DAB+ Substrate Chromogen System (#K3468, Santa Clara, CA, USA) for 3 min. The sections were counterstained with 0.1% hematoxylin. The NFIX expression was evaluated as either positive or negative across all slides. When >10% of the tumor cell nuclei were stained, it was considered positive for NFIX. Two surgical pathologists (A.I. and N.K.) followed this classification system and independently reviewed the immunoreactivity of each specimen.

### Kaplan–Meier analysis

Kaplan–Meier analysis was performed for NFIX using Kaplan–Meier Plotter software from a database of public Pan-cancer RNA-seq datasets (http://kmplot.com/analysis).

Data from 631 patients with GC were collected from a database (24). To analyze the prognostic value, the samples were divided into two groups based on the cutoff value determined by the automated software program. Hazard ratios (HRs) and P values (log-rank P) were determined for each survival analysis.

### RNA interference

Small interfering RNA (siRNA) targeting NFIX and negative control oligonucleotides were purchased from Invitrogen (#1299003, Carlsbad, CA, USA). Three independent NFIX-specific siRNAs were used in this study. Transfection of GC cell lines was conducted using Lipofectamine RNAiMAX (#13778075, Invitrogen, Waltham, Massachusetts, USA) following previously described methods (25). Specifically, 60 pmol of siRNA and 10 µL of Lipofectamine RNAiMAX were combined in 1 mL of DMEM medium to achieve a final siRNA concentration of 10 nmol/L. After incubation for 20 min, the mixture was added to the cells, which were then plated onto culture dishes. Subsequently, GC cells were analyzed 48 h after transfection, as described below.

### Quantitative Reverse Transcription-Polymerase Chain Reaction (qRT-PCR)

Total RNA was extracted using the RNeasy Mini Kit (#74004, Qiagen, Valencia, CA, USA), and 1 µg of total RNA was converted to cDNA using the First Strand cDNA Synthesis Kit (#6110A, Amersham Biosciences, Piscataway, NJ, USA). qRT-PCR was performed using an ABI PRISM 7700 Sequence Detection System (Applied Biosystems, Foster City, CA, USA). The SYBR Select Master Mix (#4472903; Applied Biosystems) was used. The primer sequences were as follows: NFIX -F CATCAAACCACTGCCCAACG, -R CTCACCAGCTCCGTCACATT; Ras association domain family member 2 (RASSF2) -F CTCTGAAGCCCCTGACTGTG, -R GTCTGTGGAGCTTGGCATCT; microRNA 21 (MIR21) -F TGACTGTTGAATCTCATGGCA, -R GTCAGACAGCCCATCGACTG; adrenoceptor alpha 2A (ADRA2A) -F CTTCTGGTTCGGCTACTGCA, -R GGCGGAAATCGTGGTTGAAG; STE20 related adaptor beta (STRADB) -F CTAGTGACCCTCTCTGGCCT, -R CACAGCCCTATGCCTCTGTC; β-actin-F CGGGAGAAATTGCAGGAGGA, and -R AAGGTCAAGACGTGCCAGAG.

### Western blot analysis

The cells were lysed as previously described (6). The lysates (30 µg) were solubilized in Laemmli sample buffer by boiling and then subjected to 10% sodium dodecyl sulfate-polyacrylamide gel electrophoresis. Following electrophoresis, the proteins were transferred onto nitrocellulose membranes and incubated with primary antibodies.

Peroxidase-conjugated IgG was used for secondary reactions. The immunocomplexes were visualized using an ECL Western Blot Detection system (RPN2235; Amersham Biosciences Corp., Piscataway, NJ, USA). Anti-β-actin (A5441, Sigma-Aldrich, St. Louis, MO, USA) was used as the loading control.

### Cell growth assay

The 3-(4,5-dimethylthiazol-2-yl)-2,5-diphenyltetrazolium bromide (MTT) assay was performed to examine cell growth as previously described (6). GC cells were seeded at 4,000 cells/well in 96-well plates. Cell growth was monitored after 1, 2, and 4 days. Three separate MTT experiments were performed, and the mean ± standard deviation (SD) was calculated.

### Spheroid colony formation assay

For spheroid generation, 1,000 cells were seeded in six-well ultra-low attachment plates (#3471, Corning, Arizona, United States). The cells were grown in mTeSR medium (#85850; STEMCELL Technologies Inc., Vancouver, BC, Canada). The plates were incubated at 37 °C in an incubator with a 5% CO2 atmosphere for 15 days. Spheroid number and size were determined using a microscope, as previously reported (25).

### Statistical analysis

The Fisher’s exact test analyzed correlations between clinicopathological parameters and NFIX expression. Significant differences between survival curves were determined using the log-rank test. Univariate and multivariate Cox regression analyses were performed to evaluate the association between clinical covariates and overall survival (OS). HRs and 95% confidence intervals (CIs) were calculated using Cox proportional hazards model. Statistical significance was set at P < 0.05 as previously (26).

## RESULTS

### Spatial transcriptome analysis enables classification of GC tissue by gene expression profile

First, we conducted a spatial transcriptome analysis. Following data quality control (QC), 2540 samples were retained for gene profile analyses. The CytAssist Visium platform enabled spatial gene expression analysis, revealing eight distinct clusters within the GC tissue. Clusters 0 and 6 represented the cancerous portions of the GC tissue, clusters 1 and 2 corresponded to the stromal compartment; cluster 3 was associated with necrotic areas; cluster 4 contained vascular structures; and cluster 5 contained smooth muscles. The distribution of these clusters and their spatial localization are depicted in the UMAP (Figure 1A) and spatial plots (Figure 1B), with representative images of each cluster (Figures 1C-1G). These observations underscore the capability of spatial transcriptome analysis to categorize GC tissues into neoplastic and non-neoplastic components based on morphological and gene expression profiles.

**Figure 1.**
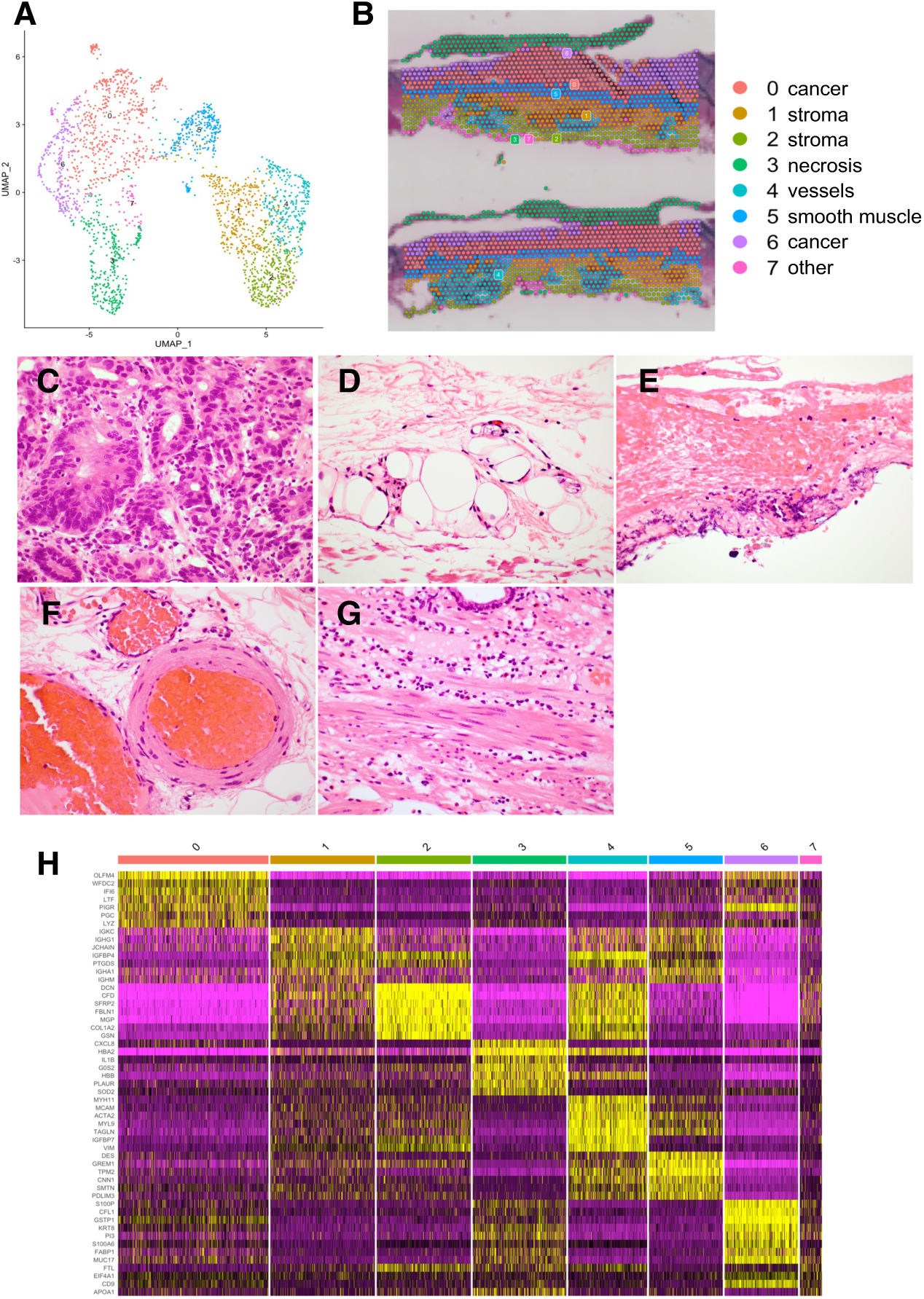
Classification of gastric cancer (GC) tissues using spatial transcriptome analysis. (A) UMAP projection of Visium (CytAssist Visium) spots also identifies 8 clusters based on differential gene expression analysis. (B) Spatial gene expression analysis classifies the cells in GC. (C-G) Representative images of GC tissue of HE for each classified cluster. (C) Cluster 0 and 6; cancer. (D) Cluster 1; stroma. (E) Cluster 3; necrosis. (F) Cluster 2; stroma. (G) Cluster 5; smooth muscle. (H) Heat map analysis of high-expression genes of each cluster.

### Identification of cancer stem cell-like cluster through spatial transcriptomic analysis

Focusing exclusively on cancer cells, we isolated clusters 0 and 6, which were identified as malignant based on their morphological characteristics, yielding 823 samples for further analysis. These samples are illustrated in UMAP (Figure 2A) and spatial plots (Figure 2B). This refined analysis revealed four sub-clusters within the cancer cell population based on their gene expression profiles (Figure 2C). These sub-clusters highlighted regions of pronounced nuclear atypia (Figure 2D), diffuse morphological characteristics predominantly located in basal areas (Figure 2E), and gland-forming components (Figure 2F).

**Figure 2.**
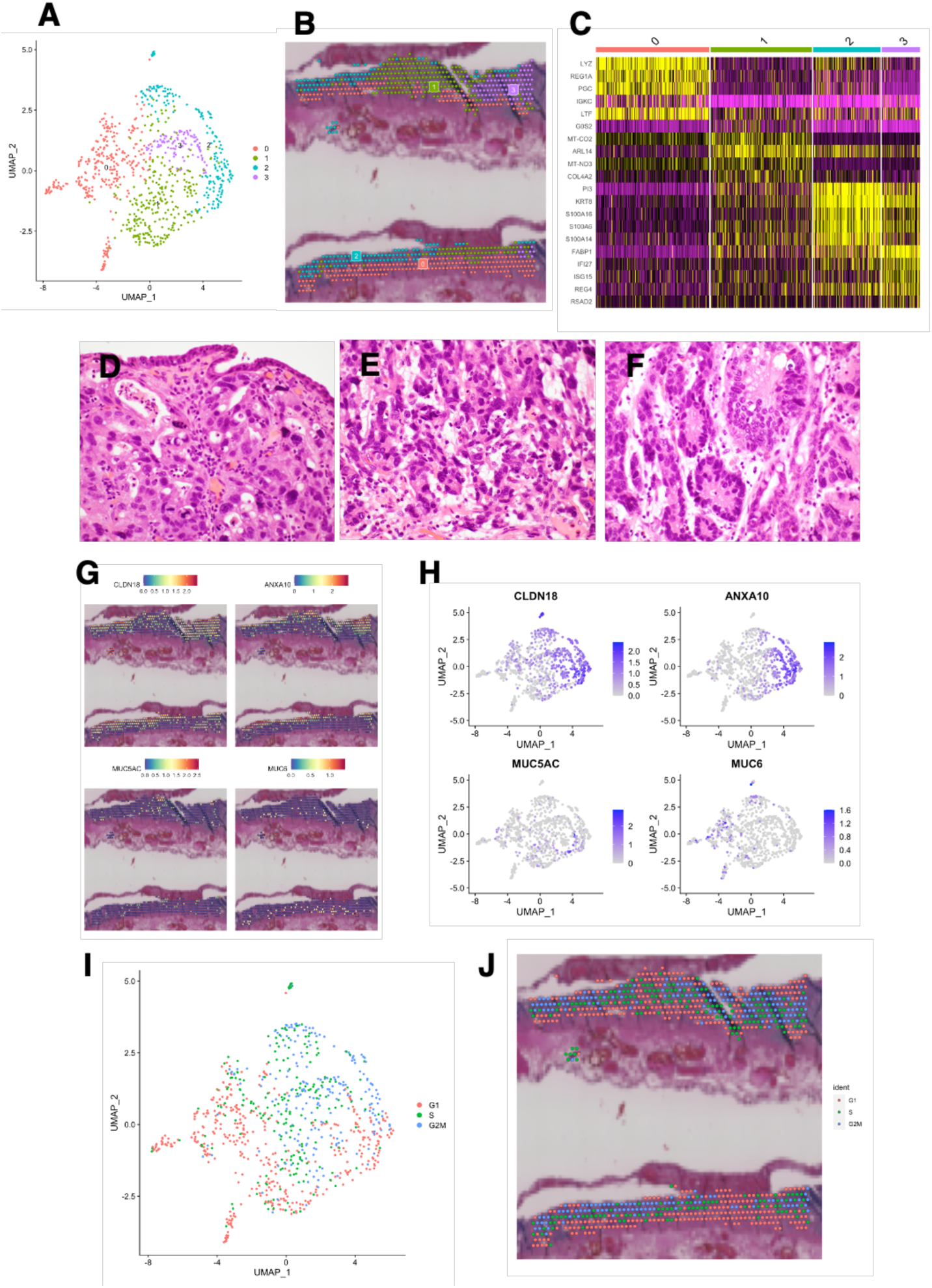
Identification of gastric cancer (GC) stem cell-like cells via spatial transcriptomic analysis. (A) UMAP projection spots identifies 4 clusters based on differential gene expression analysis of GC. (B) Spatial gene expression analysis classifies the cells in GC. (C) Heat map analysis of highly expressed genes in each cluster. (D-F) Representative images of GC of HE for each classified cluster. (D) Cluster 2; pronounced nuclear atypia area. (E) Cluster 1; cancer cells with diffuse morphological characteristics predominantly located in basal areas. (F) Cluster 3; gland-forming components. (G) Spatial analysis of CLDN18, ANXA10, MUC5AC and MUC6. (H) UMAP analysis of CLDN18, ANXA10, MUC5AC and MUC6. (I and J) Cell cycle analysis of (I) UMAP and (J) spatial plot.

Examination of specific gene expression revealed a superficial predominance of CLDN18, ANXA10, and MUC5ACdifferentiation markers, whereas MUC6, which is associated with a stem-like phenotype, was expressed in the basal layers (Figure 2G). UMAP analysis further distinguished differentiated cancer cells in clusters 2 and 3, located on the right, from stem cell-like clusters in cluster 0, which expressed MUC6 (Figure 2H). The CellCycleScoring function of the Seurat package facilitated cell cycle scoring and regression analysis, indicating G1 phase gene expression in proximity to Cluster 0, which exhibited a stem cell-like phenotype (Figures 2I and 2J). These findings suggest the presence of poorly differentiated cancer cells in cluster 0, as characterized by both morphological and genetic profiles.

### Trajectory analysis reveals a cancer stem-like cluster

Using CytoTRACE, we assessed stemness across different regions and discovered a gradient of differentiation, evidenced by lower differentiation at higher positions and greater differentiation at lower positions, which correlated with the degree of cell differentiation (Figure 3A). Gene expression analysis in relation to this gradient demonstrated elevated expression of ANXA10, CLDN18, and MUC5AC in more differentiated zones, whereas MUC6 was predominantly expressed in less differentiated cells (Figure 3B). Subsequent pseudo-time analysis (Figure 3C), with trajectories highlighted in red (Figure 3D), spatially mapped these scores (Figure 3E), with trajectories accentuated in yellow (Figure 3F). These results indicate that the cells in cluster 0, as classified by UMAP dimension reduction, exhibit less differentiation and possess more primitive characteristics according to pseudotime analysis.

**Figure 3.**
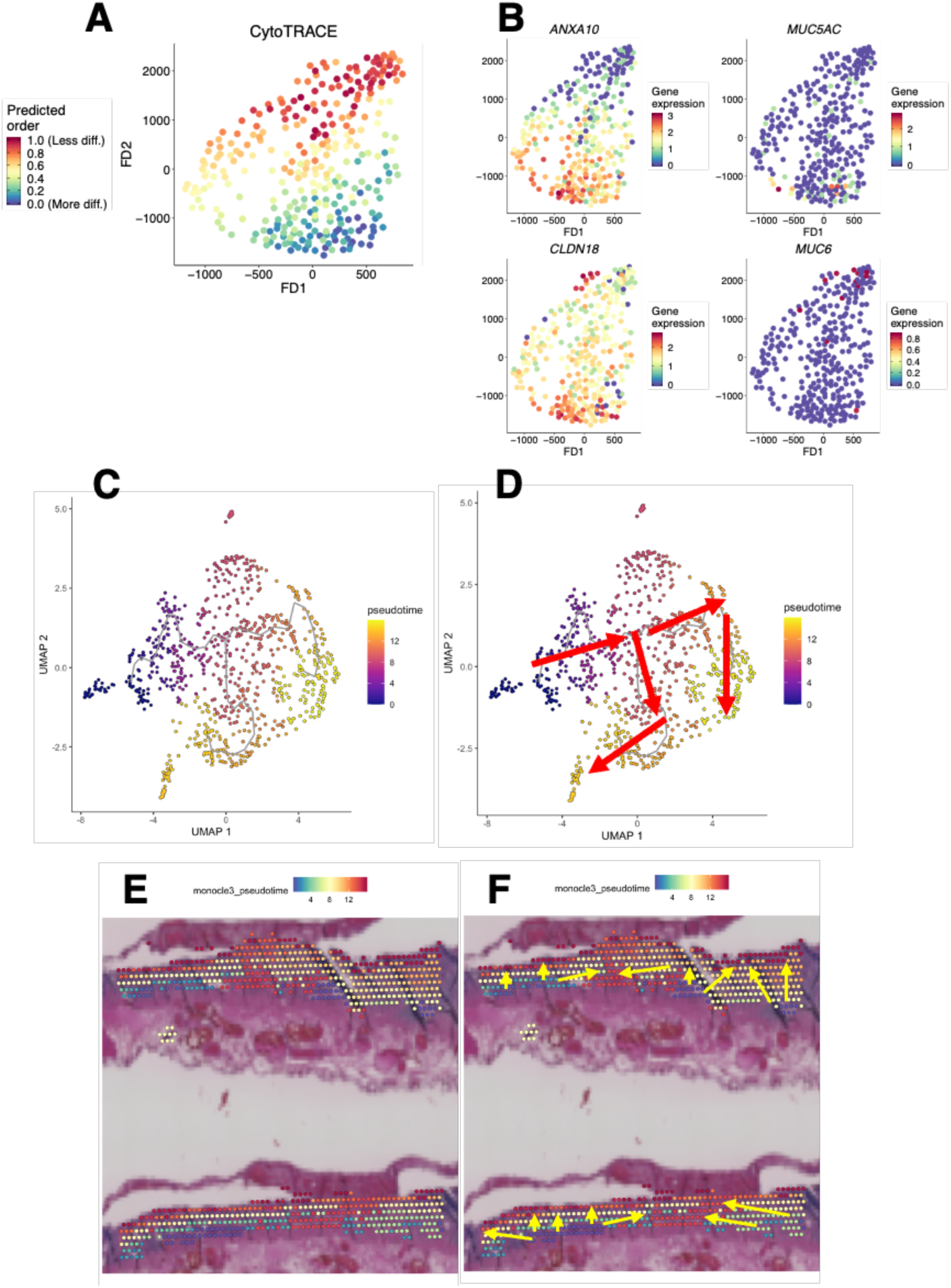
Narrowing down gastric cancer stem cell-like cells through trajectory analysis. (A and B) CytoTRACE analysis with (A) FD for dimensionality reduction and (B) gene expression distribution. (C-F) Pseudotime analysis of Monocle3. (C) UMAP and (D) flow highlighted with red arrows. (E) spatial plot and (F) flow highlighted with yellow arrows.

### Narrowing down potential target genes to identify NFIX

Figure 4A illustrates the enrichment analysis of the top 50 genes that exhibited significantly higher expression in cluster 0 compared to other clusters. This analysis revealed significant variations in genes related to glycosaminoglycan binding and peptidase regulator activity. Through the integration of morphological analysis of cluster 0 with hallmark genes derived from pseudotime and CytoTRACE analyses, we narrowed down the list of candidate genes to 16. Their expression was then demonstrated in UMAP (Figure 4C). Among these, NFIX has emerged as a novel target in GC, showing predominantly basolateral expression (Figure 4D). This comprehensive spatial transcriptome analysis identified NFIX as a pivotal gene linked to stem cell-like cells in GC. Further investigation into its potential as a therapeutic target for GC is warranted.

**Figure 4.**
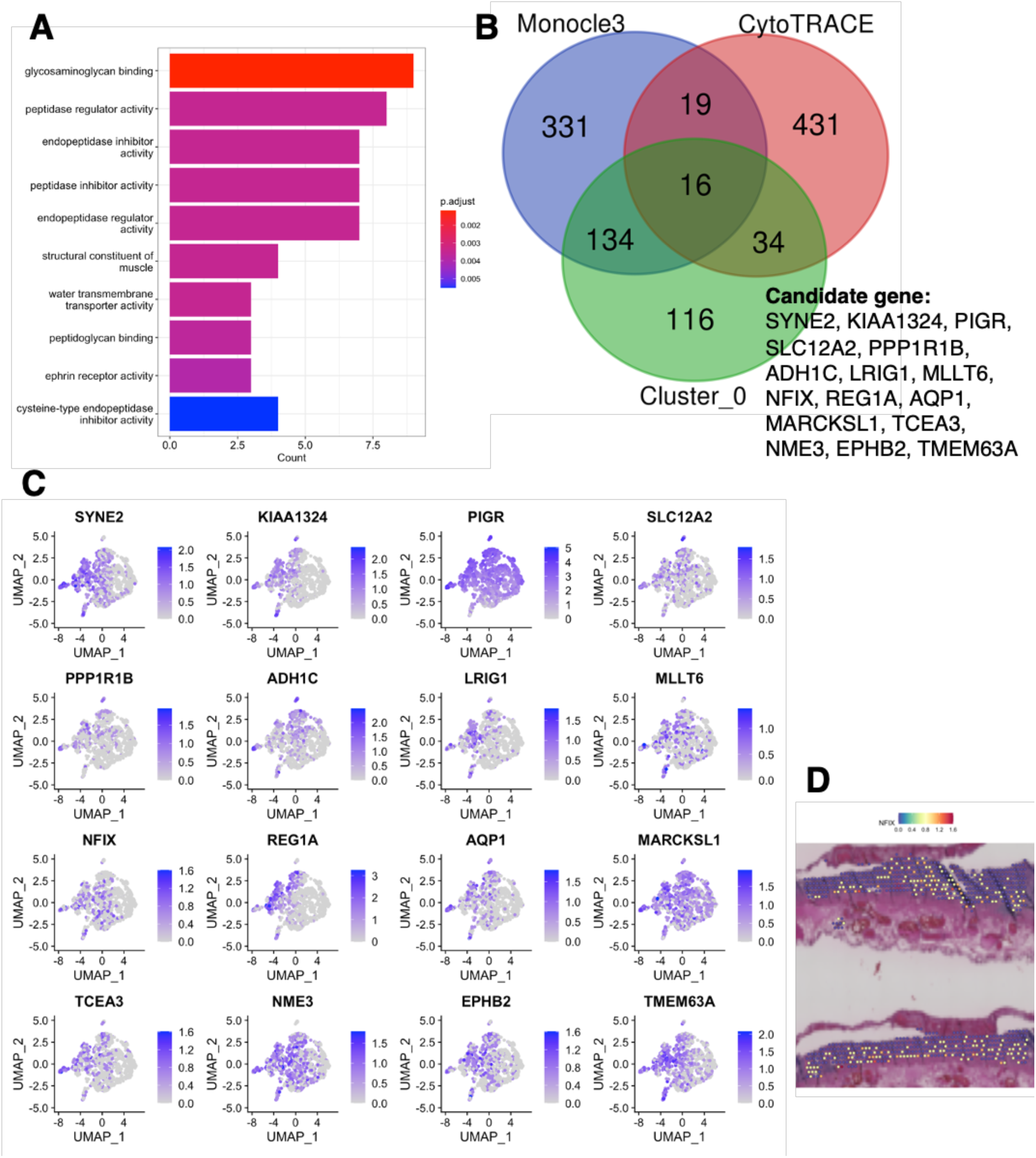
Identification of NFIX as a key gene associated with cancer stemness in gastric cancer. (A) Gene Ontology enrichment analysis of the top 50 genes of cancer stem cell-like cells. (B) The overlap of the Venn diagram showed that there are 16 candidate targeted genes predicted by Cluster 0, Monocle3, and CytoTRACE. (C) Feature plots of each candidate gene. (D) Spatial plot of NFIX.

### NFIX expression and clinico-pathologic characteristics in GC

Next, we focused our analysis on NFIX among the candidate genes because NFIX has not been previously studied immunohistochemically in GC. We observed NFIX expression in stomach tissue. Immunohistochemical staining was performed on specimens from 127 GC patients to confirm the distribution of KIF18B expression. In the non-neoplastic stomach, NFIX expression was observed in the isthmus (Fig. 5a and 5b), a region containing somatic stem cells. NFIX is not expressed in non-neoplastic gastric mucosa, including intestinal metaplasia. NFIX was considered positive when NFIX expression was detected in more than 10% of the tumor cells. Accordingly, immunohistochemical staining of 127 GC samples was performed, of which 65 (51%) were found positive for NFIX. Based on these results, we examined the correlation between NFIX expression and various clinicopathological factors (Table 1). pT stage (p = 0.035), pM stage (p < 0.01), and stage (p = 0.023) were correlated with NFIX expression. These results indicated that NFIX was expressed in isthmus and 51 % GC cases and that its expression correlates with pT stage, pM stage, and stage.

**Figure 5.**
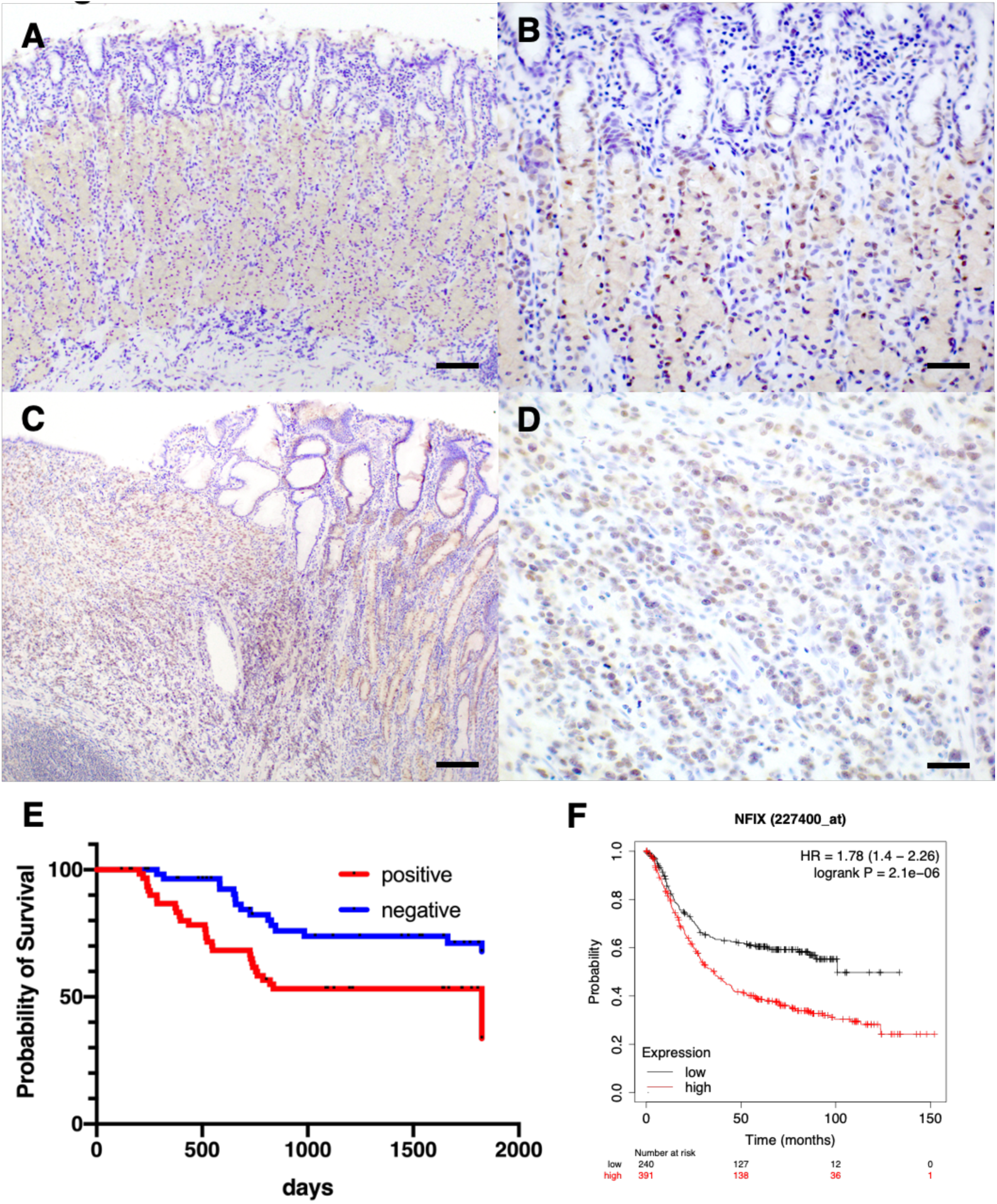
Correlation of NFIX expression with poor prognosis in gastric cancer (GC). (A-D) Representative immunohistochemical images of NFIX. (A and B) Immunohistochemical staining of NFIX in non-neoplastic gastric mucosa. Original magnification: (A) 100×; scale bars, 200 µm and (B) 400×; scale bars, 50 µm. (C and D) Immunohistochemical staining of NFIX in GC. Original magnification: (C) 100×; scale bars, 200 µm and (D) 400×; scale bars, 50 µm. (E) Overall survival probability in the 127 GC cases. (F) Overall survival probability in the public RNA-seq dataset.

**Table 1.** Relationship between nuclear factor I X (NFIX) expression and clinicopathologic characteristics of the 127 gastric cancer cases.

|  |  | NFIX expression |  | *p* Value |
| --- | --- | --- | --- | --- |
|  |  | Positive (%) | Negative |  |
| Age (years) |  |  |  | 0.723 |
|  | <71 | 33 (49) | 34 |  |
|  | ≥71 | 32 (53) | 28 |  |
| Sex |  |  |  | 0.239 |
|  | Male | 44 (48) | 48 |  |
|  | Female | 21 (60) | 14 |  |
| pT stage |  |  |  | 0.035 |
|  | pT1/2 | 40 (45) | 49 |  |
|  | pT3/4 | 25 (66) | 13 |  |
| pN stage |  |  |  | 0.258 |
|  | pN0 | 18 (43) | 24 |  |
|  | pN1/2/3 | 47 (55) | 38 |  |
| pM stage |  |  |  | < 0.01 |
|  | pM0 | 54 (47) | 61 |  |
|  | pM1 | 11 (92) | 1 |  |
| Stage |  |  |  | 0.023 |
|  | I / II | 18 (40) | 27 |  |
|  | III / IV | 47 (57) | 35 |  |
| Histology |  |  |  | 0.067 |
|  | Intestinal | 15 (36) | 27 |  |
|  | Diffuse | 50 (59) | 35 |  |

### Correlation between NFIX expression and TCGA-like molecular subtypes

Next, we analyzed the molecular subtypes that showed marked histological diversity. The 127 GC cases were classified into four molecular subtypes based on the expression patterns of surrogate markers: 10 cases (7.9 %) of EBV-like, 22 cases (17.3 %) of MSI-like, 59 cases (46.5 %) of GS-like, and 51 cases (28.3 %) of CIN-like subtype (Table 2). NFIX expression significantly correlated with the GS-like subtype (p < 0.01) and significantly inversely correlated with the MSI-like subtype (p < 0.01) (Table 2).

**Table 2.**
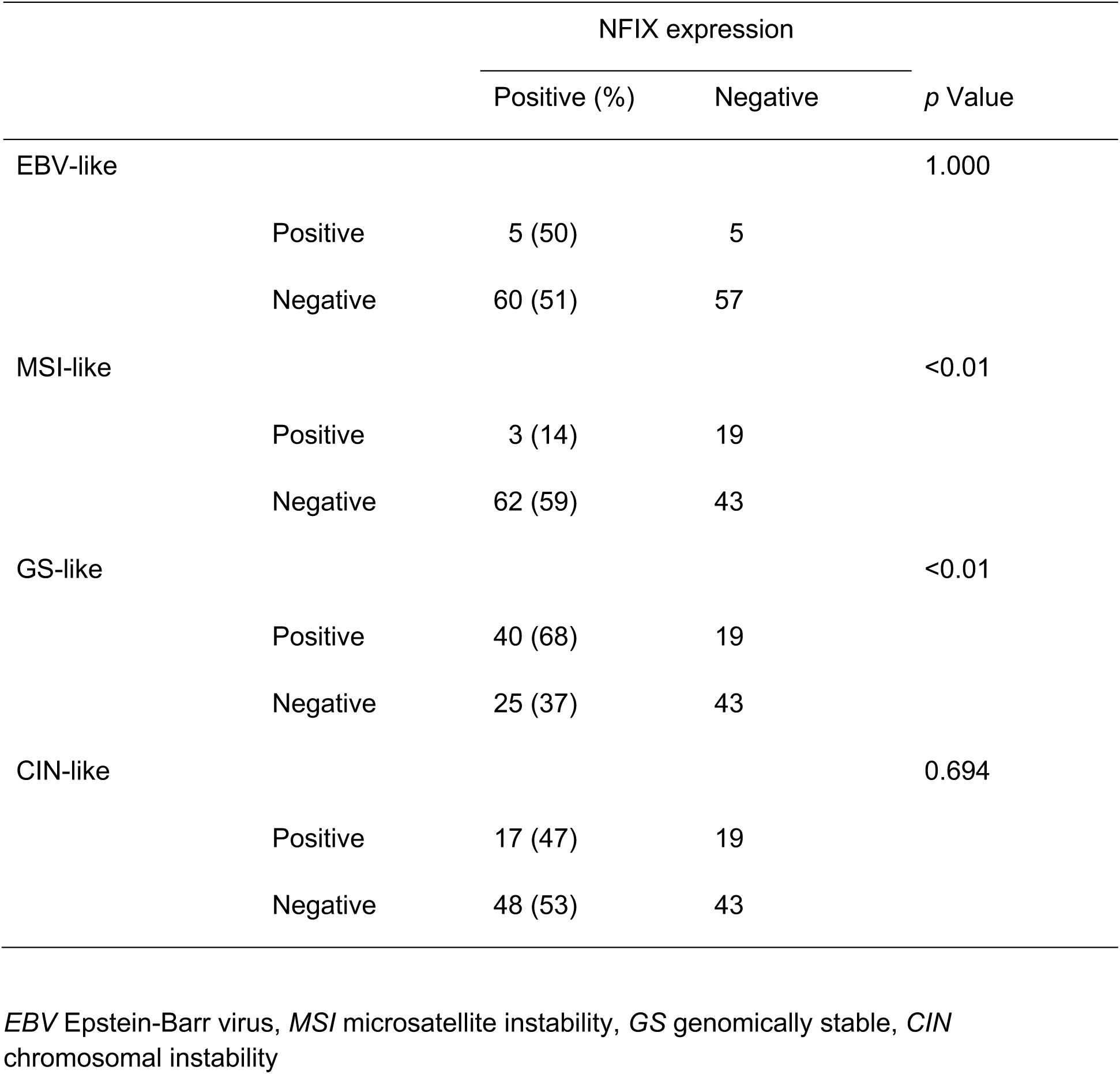
The relationship between nuclear factor I X (NFIX) expression and the Cancer Genome Atlas (TCGA)-like classification in the 127 gastric cancer cases.

### NFIX expression and survival analysis in GC

Kaplan–Meier curves were drawn, and log-rank tests were performed to determine whether NFIX expression in GC affected prognosis. NFIX-negative GC cases showed a significantly better OS probability than NFIX-negative GC cases (P < 0.01; Fig. 5E). To confirm this result, we also assessed a public dataset that included 631 GC cases, and similar results were obtained, with reduced NFIX expression of NFIX significantly associated with a poorer prognosis (P < 0.01; Fig. 5F). As we found that NFIX expression was significantly associated with poor prognosis, we performed univariate and multivariate analyses using Cox hazard analyses (Table 3). Univariate analysis revealed several prognostic factors, including NFIX expression, pT stage, pN stage, pM stage, and NFIX expression (P < 0.01). Multivariate analysis revealed that pT, pN, and pM stages were independent prognostic factors in patients with GC. NFIX expression was not an independent factor for poor prognosis (HR, 1.480; 95% CI, 0.761-2.880; P = 0.248). These results indicate that although NFIX expression is associated with poor prognosis, it is not an independent poor prognostic factor.

**Table 3.** Univariate and multivariate Cox regression analyses of nuclear factor I X expression and survival in the 127 gastric cancer cases.

| Features | Univariate analysis |  | Multivariate analysis |  |
| --- | --- | --- | --- | --- |
|  | HR (95%CI) | *p* Value | HR (95%CI) | *p* Value |
| Age |  | 0.230 |  |  |
| ≥ 71 | 1 (ref.) |  |  |  |
| < 71 | 1.408 (0.806-2.461) |  |  |  |
| Sex |  | 0.376 |  |  |
| Female | 1 (ref.) |  |  |  |
| Male | 1.308 (0.722-2.371) |  |  |  |
| pT stage |  | < 0.01 |  | 0.033 |
| pT1/2 | 1 (ref.) |  | 1 (ref.) |  |
| pT3/4 | 3.000 (1.716-5.230) |  | 1.915 (1.054-3.480) |  |
| pN stage |  | < 0.01 |  | <0.01 |
| pN0 | 1 (ref.) |  | 1 (ref.) |  |
| pN1/2/3 | 7.565 (2.355-24.31) |  | 6.052 (1.869-19.60) |  |
| pM stage |  | < 0.01 |  | 0.042 |
| pM0 | 1 (ref.) |  | 1 (ref.) |  |
| pM1 | 3.514 (1.836-6.730) |  | 2.078 (1.026-4.210) |  |
| NFIX expression |  | <0.01 |  | 0.248 |
| Negative | 1 (ref.) |  | 1 (ref.) |  |
| Positive | 2.534 (1.384-4.641) |  | 1.480 (0.761-2.880) |  |

#### NFIX expression acts in a tumor-promoting way by regulating kinase activity

Next, we tested whether the functions of NFIX discovered thus far were also found in human-derived GC cell lines. Western blot analysis revealed NFIX expression in HFE- 145 and MKN-45 cells (Fig. 6A). NFIX knockdown was validated using an NFIX-specific siRNA in MKN-45 (Fig. 6B) and HFE-145 cells (Supplementary Fig. 1A). The proliferation assay revealed that the growth of these NFIX siRNA-transfected cells were significantly lower compared to that of the negative control siRNA-transfected cell lines (Fig. 6C). We examined the migration activity of NFIX siRNA-transfected cells using a wound healing assay. In this assay, the wound-healing distance of NFIX siRNA-transfected cells was significantly lower than that of negative control siRNA-transfected MKN-45 cells (Fig. 6D and 6E). We analyzed the association between NFIX expression and spheroid colony formation in the GC cell lines. In MKN-45 cells, both the number and size of spheroids significantly decreased in NFIX siRNA-transfected cells (Fig. 6F). These assays were also performed on a non-tumor gastric epithelial cell line, HFE-145, with similar results as the proliferation (Supplementary Fig. 1B), wound healing (Supplementary Fig. 1C and 1D), and spheroid colony formation assays (Supplementary Fig. 1E).

**Figure 6.**
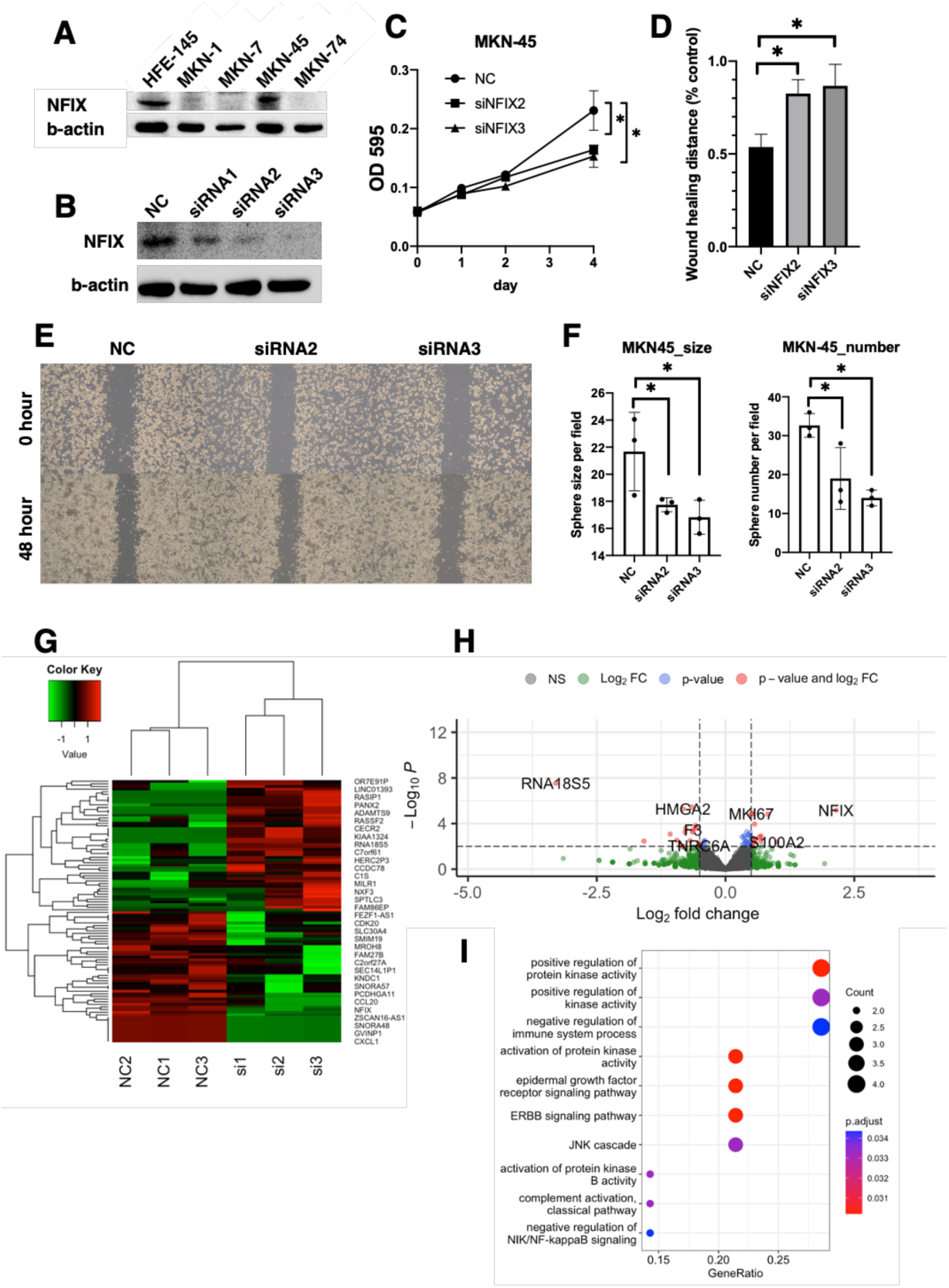
Role of NFIX in gastric cancer (GC) cells: proliferation, stemness, and kinase activity regulation. (A) Western blot analysis of NFIX in five non-neoplastic or GC cell lines. (B) Western blot analysis of NFIX in MKN-45 cells transfected with the negative control or NFIX siRNA. (C) Effect of NFIX knockdown on cell growth in MKN-45 cells transfected with the negative control or NFIX siRNA. (D and E) Wound-healing assay in MKN-45 cells transfected with the negative control or NFIX siRNA. (D) The mean percentage of wound closure. (E) Representative image. (F) Number and size of spheroids formed by MKN-45 cell lines transfected with negative control or NFIX siRNA. (G) The heatmap shows the gene expression profile of 3 independent sets of MKN-45 cells transfected with the negative control or NFIX siRNA by RNA sequencing analysis. Data are color-coded to reflect the gene expression level. (H) Volcano plot comparing MKN-45 cells transfected with the negative control and that with NFIX siRNA. (I) Gene Ontology enrichment analysis of the top 50 genes.

We then performed a bioinformatics study of the functions based on changes in gene profiles upon the knockdown of NFIX. Upon comparison between NFIX siRNA-transfected cells and negative control siRNA-transfected cells, a total of 99 DEGs were identified. Among these, 61 were found to be upregulated while 38 were downregulated. The heatmap shows representative DEGs (Fig. 6G), and the volcano plot shows the distribution of all DEGs in the dimensions of -log (FDR) and log FC (Fig. 6H). Gene ontology (GO) analysis was conducted utilizing the top 20 genes expressed in the NFIX- expressing state, specifically in the negative control siRNA-transfected cells (Fig. 6I). "Positive regulation of protein kinase activity" and other genes that control kinase activity were found to vary (Supplementary Fig. 2A). Therefore, we further examined whether this could be confirmed by other means, such as qRT-PCR, to confirm that gene expression, such as that of NFIX, was as variable as observed by RNA sequencing (Supplementary Fig. 2B). These results suggest that NFIX promotes tumor growth by regulating kinase activity.

## DISCUSSION

In this study, we advanced the classification of GC tissues through spatial transcriptome analysis by identifying candidate genes with therapeutic potential. Our focused investigation of NFIX, guided by both morphological assessments and bioinformatic insights, is pivotal. Through comprehensive examination across our cohort, we underscored the critical role of NFIX in the pathogenesis and progression of GC.

In our study, 10x Visium CytAssist system (27) was employed for spatial transcriptome analysis of GC, which is a novel approach given the rarity of such analyses in this field (13, 28) and the recent development of this technology. This study represents the inaugural application of the CytAssist system in GC, marking a significant advancement in our understanding of the spatial biology of GC. Moreover, while previous studies have explored stemness within a non-neoplastic gastric context (15, 29), our investigation uniquely focused on CSC-like aspects, setting a precedent in the exploration of GC. We have refined our approach to identifying stemness not merely through morphological classification but by integrating gene expression profiles with differentiation markers, such as ANXA10 (9, 30) and MUC5AC (31). This methodological rigor, coupled with trajectory analysis, facilitated the identification and clustering of stem cell-like cells within GC tissue. Through these comprehensive analyses, we successfully identified NFIX as a critical factor underscoring its potential as a therapeutic target.

This study represents a pioneering effort to elucidate the role of NFIX in GC through immunohistochemical analysis and is the first report of its kind. Our findings revealed that NFIX plays a crucial role in modulating cancer stemness and kinase activity during GC progression. NFIX belongs to the NFI family, which also includes NFIX, NFIB, and NFIC. NFIX contributes to muscle development, muscular dystrophy, hematopoiesis, cancer, and neural stem cell biology (32). NFIX has been reported to be attenuated in medulloblastoma (33) and colorectal cancer (34) and acts on miRNAs. NFIX has been correlated with pT and pM stages in our cohort studies and is involved in proliferation and colony-forming ability in cell line experiments, successfully supporting this clinical attitude. Furthermore, in the study on the correlation between TCGA classification and NFIX, NFIX expression was correlated with GS-like type GC and inversely correlated with MSI-like type GC. This suggests that MSI-type GC may benefit from treatment with immune checkpoint inhibitors (35).Conversely, discovery of molecules characteristic of GS-type gastric cancer may revolutionize the treatment of gastric cancer. By uncovering the multifaceted involvement of NFIX in GC, our research not only advances the current understanding of the molecular underpinnings of this malignancy, but also highlights NFIX as a potential biomarker and therapeutic target, offering new avenues for targeted interventions in GC treatment.

This study had certain limitations that warrant further discussion. First, spatial transcriptome analysis was conducted for a single case. Although this may appear to be a significant constraint, it is mitigated by subsequent validations. The relevance of the identified molecule was robustly confirmed through immunostaining in our cohort and reinforced by extensive reviews of public databases, thereby substantiating our findings despite the initial limitations. Second, the resolution of spatial transcriptome analysis poses another limitation. The system employed in our study offers a resolution of approximately 90 micrometers (11, 12), precluding single-cell-level analysis. Despite this limitation, the utility of this method for identifying potential therapeutic target genes remains to be validated through supplementary approaches. This ensured that the resolution of the method, which was not ideal for single-cell analysis, was sufficiently robust to satisfy the objectives of our study. Finally, the retrospective nature of our study represents a limitation, as it was not a prospective study. Implementation of in vivo studies and prospective clinical trials is essential for further validation of our findings. Such studies would not only corroborate our results but also enhance the clinical applicability of our findings.

In conclusion, our application of spatial transcriptome analysis significantly advanced the precision in classifying GC tissues, facilitating a refined approach for identifying candidate genes for therapeutic intervention. Specifically, this study highlighted NFIX as a pivotal gene of interest. The expression of NFIX not only correlates with the prognosis of GC but also plays a critical role in disease progression through its kinase activity. These findings highlight the potential of NFIX as a novel therapeutic target and offer promising avenues for tailoring treatment strategies for GC. Through this research, we contribute to the ongoing efforts to enhance patient outcomes by elucidating the molecular underpinnings of gastric cancer and opening new pathways for targeted therapies.

## ACKNOWLEDGEMENTS

The authors are grateful to the patient for providing consent to report their clinical information and data. We thank R. Nobuhiro, S. Norimura, and Y. Kurokawa for their technical assistance in this study. We thank the NGS Core Facility of the Research Institute for Microbial Diseases of Osaka University.

## DISCLOSURE

### Funding Information

This study was supported by a KAKENHI Grant-in-Aid for Early-Career Scientists (Grant Number: JP21K15394) from the Japan Society for the Promotion of Science.

### Conflict of Interest

The authors have no conflict of interest.

### Ethics approval and consent to participate

This study was approved by the Ethics Committee for Human Genome Research of Hiroshima University (E2001-9923), and was conducted in accordance with the guidelines of the Declaration of Helsinki. The opt-out method was used in this retrospective study. Written informed consent was obtained from all patients. An opt-out informed consent protocol was used. This procedure was reviewed and approved by the Ethics Committee for Human Genome Research of Hiroshima University (approval number E2001-9923, date of decision: April 5, 2023).

### Author Contributions

AI designed this study. KK, TS, SO, and TK collected and analyzed the patients’ clinical data. AI and NU performed the experiments and collected and analyzed the data. AI, TF, AK, NK, KS, and NO interpreted and analyzed the results. HA and DS were used to generate the HFE-145 cells. AI and WY drafted and edited the manuscript. All authors have read and approved the manuscript and agree to be accountable for all aspects of the research to ensure that the accuracy or integrity of any part of the work is appropriately investigated and resolved.

---

**Supplementary Figure 1.**
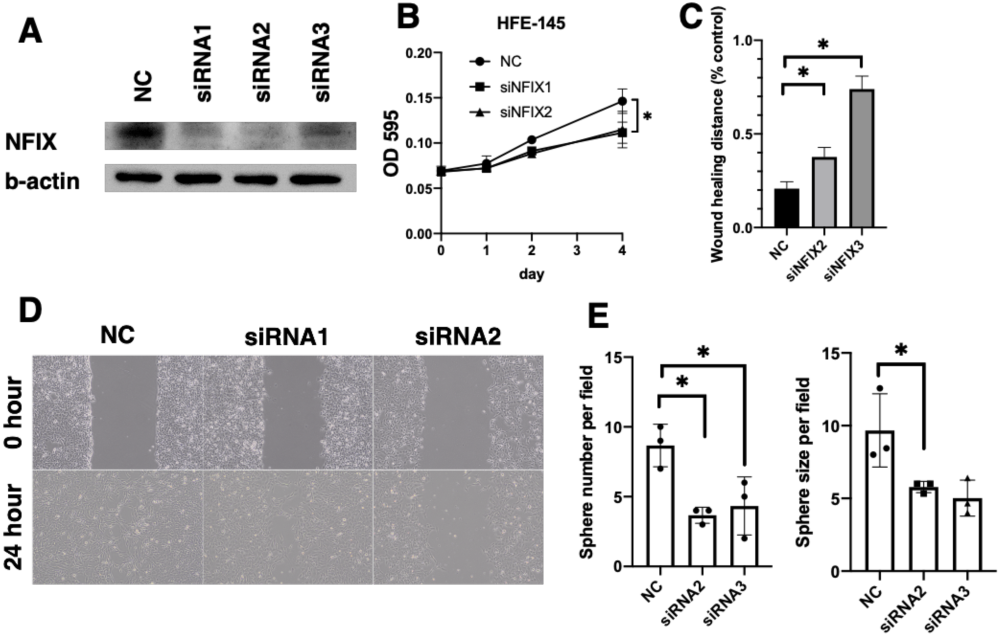
Proliferative capacity and stemness in non-gastric cancer cell lines. (A) Western blot analysis of NFIX in HFE-145 cells transfected with the negative control or NFIX siRNA. (C) Effect of NFIX knockdown on cell growth in HFE-145 cells transfected with the negative control or NFIX siRNA. (C and D) Wound-healing assay in HFE-145 cells transfected with the negative control or NFIX siRNA. (C) The mean percentage of wound closure. (D) Representative image. (E) Number and size of spheroids formed by MKN-45 cell lines transfected with negative control or NFIX siRNA.

**Supplementary Figure 2.**
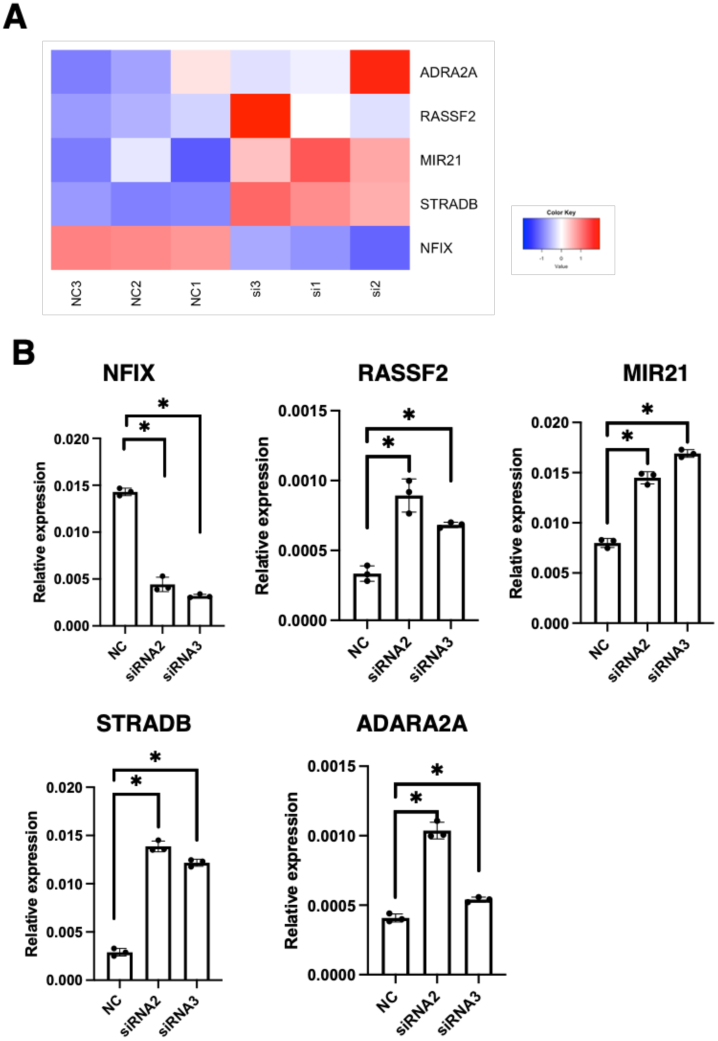
Confirmation of genetic variation using gastric cancer cell lines. (A) Heatmap of genes found to be related in RNA sequence. Quantitative reverse transcription-polymerase chain reaction analysis of NFIX, RASSF2, MIR21, STRADB and ADARA2A genes in MKN-45 cells transfected with the negative control or NFIX siRNA.

## Notes

### Competing Interest Statement

The authors have declared no competing interest.

